# Novel canine high-quality metagenome-assembled genomes, prophages, and host-associated plasmids by long-read metagenomics together with Hi-C proximity ligation

**DOI:** 10.1101/2021.07.02.450895

**Authors:** Anna Cuscó, Daniel Pérez, Joaquim Viñes, Norma Fàbregas, Olga Francino

## Abstract

Long-read metagenomics facilitates the assembly of high-quality metagenome-assembled genomes (HQ MAGs) out of complex microbiomes. It provides highly contiguous assemblies by spanning repetitive regions, complete ribosomal genes, and mobile genetic elements. Hi-C proximity ligation data bins the long contigs and their associated extra-chromosomal elements to their bacterial host. Here, we characterized a canine fecal sample combining a long-read metagenomics assembly with Hi-C data, and further correcting frameshift errors.

We retrieved 27 HQ MAGs and seven medium-quality (MQ) MAGs considering MIMAG criteria. All the long-read canine MAGs improved previous short-read MAGs from public datasets regarding contiguity of the assembly, presence, and completeness of the ribosomal operons, and presence of canonical tRNAs. This trend was also observed when comparing to representative genomes from a pure culture (short-read assemblies). Moreover, Hi-C data linked six potential plasmids to their bacterial hosts. Finally, we identified 51 bacteriophages integrated into their bacterial host, providing novel host information for eight viral clusters that included Gut Phage Database viral genomes. Even though three viral clusters were species-specific, most of them presented a broader host range.

In conclusion, long-read metagenomics retrieved long contigs harboring complete assembled ribosomal operons, prophages, and other mobile genetic elements. Hi-C binned together the long contigs into HQ and MQ MAGs, some of them representing closely related species. Long-read metagenomics and Hi-C proximity ligation are likely to become a comprehensive approach to HQ MAGs discovery and assignment of extra-chromosomal elements to their bacterial host.

## Background

Complex microbiomes are a source of novel bacterial diversity, but cultivation methods fail to isolate all these species. Alternatively, metagenomics provides sequence information of all the DNA from a microbiome sample and retrieves metagenome-assembled genomes (MAGs) that can represent novel uncultured bacteria [1,2].

Short-read derived MAGs are usually fragmented and lack ribosomal gene sequences, whose presence is required to be considered high-quality [3]. Ribosomal genes are the most widely used taxonomic markers to classify bacteria since they present highly conserved regions to design universal primers and hypervariable regions with taxon-specific divergences [4]. Since they are repeated and highly conserved, short-read metagenomics collapses these genes together and cannot locate them in their respective bacterial genome [5].

Long-read metagenomics uses long DNA stretches, solving many issues from short-read derived MAGs. Long-read sequencing spans complete ribosomal genes and their genomics context, bridging together microbiome insights obtained by short-read MAGs and 16S rRNA sequencing surveys [6]. Besides, it spans complete mobile genetic elements (MGE) such as prophages or plasmids [7–10] that can harbor antimicrobial resistance genes or virulence factors. Sequencing full-length MGE and locating them correctly in the chromosome or plasmid can unravel horizontal gene transfer events or the pathogenic potential of a specific microorganism [11].

However, long-read sequencing needs to overcome two main issues: obtaining long DNA fragments and reducing the sequencing error rate. For the first one, high-molecular weight DNA extractions suited for sample type work efficiently producing long-reads, as previously demonstrated for fecal samples [12]. For the second one, the higher error rate when compared to other technologies can be significantly reduced by deep sequencing [13] and by using error-specific correction software, such as frameshift-aware software for Nanopore sequencing [14].

To further disentangle complex microbiomes, metagenomics can be complemented with Hi-C proximity ligation data. Hi-C proximity ligation cross-links DNA *in vivo* within intact cells to capture interactions between DNA molecules in close physical proximity [15,16]. This approach further improves the contiguity of a metagenome assembly and captures interactions between plasmids or viruses and their host genomes. To date, only two studies have combined long-read metagenomics with Hi-C proximity ligation data: in a cow rumen, to link viruses and antimicrobial resistance genes to their microbial host [17] and in a sheep gut, to generate “lineage-resolved” MAGs [18].

This study aimed to characterize a canine fecal sample by combining the long-read assembly and Hi-C proximity ligation data to unravel high-quality MAGs and their associated extra-chromosomal elements.

## Material and methods

### Long-read metagenomics: DNA extraction and Nanopore sequencing

Our study focuses on the microbiome analysis of a single fecal sample of a healthy dog. Using the same fecal sample, we extracted High-Molecular Weight (HMW) DNA through Quick-DNA HMW MagBead (Zymo Research) and non-HMW DNA through DNA Miniprep Kit (Zymo Research). We prepared a sequencing library for each DNA extraction using the Ligation Sequencing Kit 1D (SQK-LSK109; Oxford Nanopore Technologies) and sequenced each of them in a single Flowcell R9.4.1 using MinION™ (Oxford Nanopore Technologies). More details were described previously [19].

### Hi-C metagenome cross-linking, and Illumina sequencing

The same fecal sample was used to generate a Hi-C library using the ProxiMeta Hi-C kit following the manufacturer’s protocol (Phase Genomics). The Hi-C method cross-links DNA molecules that are in close physical proximity within intact cells. Hi-C libraries were sequenced on an Illumina HiSeq 4000 platform, generating 75 bp paired-end reads.

### Metagenome assembly and deconvolution

Raw fast5 files from Nanopore sequencing were basecalled using Guppy 3.4.5 (Oxford Nanopore Technologies) with high accuracy basecalling mode (dna_r9.4.1_450bps_hac.cfg). During the basecalling, the reads with an accuracy lower than 7 were discarded.

Before proceeding with the metagenomics assembly, we performed an error-correction step of the raw Nanopore reads using canu 2.0 [20]. We merged the two Nanopore runs data and performed the metagenome assembly with Flye 2.7 [21] (options: --nano-corr --meta, --genome-size 500 m, --plasmids). We polished the Flye assembly with one round of medaka 1.0.1(https://nanoporetech.github.io/medaka/), including all the raw Nanopore fastq files as input.

We uploaded the metagenome assembly and the raw Hi-C sequencing data to the ProxiMeta cloud-based pipeline (Phase Genomics, December 2020), where it was processed, and the final metagenomics bins were retrieved.

### Characterization of the high-quality and medium-quality MAGs

We further corrected the metagenomics bins by correcting the frameshift errors, as described in [14], using Diamond 0.9.32 [22] and MEGAN-LR 6.19.1 [23]. We classified our MAGs considering MIMAG criteria [3] as high-quality MAG (HQ MAG), when is > 90% complete, and presents < 5% contamination, rRNAs genes and tRNAs; and medium-quality MAG (MQ MAG), when is > 50% complete and presents < 10% contamination.

To assess the novelty and the taxonomy of the metagenomic bins, we used GTDB-tk 1.3.0 [24] with GTDB taxonomy release 95 [25]. FastANI 1.3 [26] was used to determine the average nucleotide identity (ANI) between related genomes.

We used Prokka 1.13.4 [35] to annotate the genomes and assess the number of coding sequences (CDS), ribosomal genes, and tRNAs of the MAGs. Since the ribosomal genes are together within the *rrn* operon, when the number of 16S rRNAs, 23S rRNAs, and 5S rRNAs was not the same within a MAG, we double-checked their presence using RNAmmer 1.2 [27] server.

We compared the HQ MAGs obtained to previously reported MAGs from the most extensive and recent gastrointestinal collections: i) the animal gut metagenome [28], which includes MAGs from the dog gut catalog [29], and ii) the Unified Human Gastrointestinal Genome (UHGG) [2]. We retrieved MAGs representing the same species as our HQ MAGs by keeping: i) those with > 95% of ANI [26] for the animal gut metagenome; and ii) those with the equivalent species-level taxonomy as stated by GTDB-tk for UHGG.

Finally, we performed a pangenome analysis using Anvi’o 7 [30] for *Phocaeicola* species (includes some former *Bacteroides* species [31]). Within Anvi’o pangenomics workflow [32], Prodigal [33] was used as a gene caller to identify open reading frames (ORFs), whereas genes were functionally annotated using blastp against NCBI COGs database [34]. We created the pangenome database using NCBI’s blastp to calculate each amino acid sequence’s similarity in every genome against every other amino acid sequence across all genomes and subsequently to resolve gene clusters. We set the MCL inflation parameter to 4, and we used pyANI to calculate the Average Nucleotide Identity (ANI) values between the genomes [32].

### Plasmid analysis

We assessed the genomic bins representing HQ MAGs and MQ MAGs with < 5% contamination for any putative plasmids.

The putative plasmids within our HQ MAGs and MQ MAGs were predicted using Plasflow 1.1.0 [35]. They were further annotated with Prokka 1.14.6 [35] to identify plasmid-associated genes, and with Abricate 0.8.13 (https://github.com/tseemann/abricate) to identify potential antimicrobial-resistant genes with CARD database [36] or virulence factors with VFDB database [37].

We further inspected the putative plasmids by assessing: i) blast results against nr/nt NCBI database; ii) their relative coverage when compared to the associated bacterial host (from Flye 2.7 [21] output), and iii) and their circularity (from Flye 2.7 [21] output).

### Bacteriophage analysis

VirSorter2 2.1 [38] and Vibrant 1.2.1 [39] were used to detect viruses within the HQ MAGs and MQ MAGs. CheckV 0.7.0 (https://bitbucket.org/berkeleylab/checkv/) was used to assess single-contig viral genomes’ quality and remove potential host contamination within integrated viruses. If Virsorter2 and Vibrant redundantly detected a viral signal, we kept the one with the highest quality and completeness. We used vConTACT2 0.9.19 [40] to cluster viral sequences and provide taxonomic context. The results reported here are from high-quality and medium-quality predicted viruses. Low-quality predicted viruses were not included.

To perform vConTACT2, we used a subset of the Gut Phage database (GPD) [41]. To create this subset, we mapped our predicted bacteriophages to the whole GPD (n=142,809) using Minimap2 2.17 [42]. The GPD viral genomes that mapped with our predicted bacteriophages (n=682) and our predicted bacteriophages were included as input sequences into vConTACT2. Then we predicted the proteins using Prodigal 2.6.3 [33] and run vConTACT2 against its ProkaryoticViralRefSeq201-merged database. The resulting network was visualized using Cytoscape 3.8.2 [43].

## Results

We characterized the fecal metagenome of a healthy dog combining a long-read metagenomics assembly and Hi-C proximity ligation data. After the two Nanopore runs, we obtained a total of 16.94 million reads (36.05 Gb). The long reads were assembled using MetaFlye into a 142 Mbp metagenomics assembly, with a mean contig size of 150,083 bp. The Proximeta Hi-C library was sequenced with Illumina producing 75.01 million paired-end reads (11.40 Gb). The long-reads metagenomics assembly and the Hi-C paired-end reads were uploaded to the ProxiMeta analysis cloud to retrieve the genomic bins. We applied an experimental binning step by proximity ligation, linking contigs in close physical proximity within an intact cell. We further corrected the metagenomics bins by correcting the frameshift errors and proceeded with their characterization as detailed on Additional File 1. The highly complete genomic bins were representing high-quality (HQ) and medium-quality (MQ) MAGs that we named as CanMAGs, short for Canine MAGs.

### Long-read contigs included ribosomal genes, and Hi-C data binning retrieved HQ MAGs and detected plasmid-chromosome interactions

Combining a long-read metagenomics assembly with Hi-C proximity ligation data, followed by a frameshift-correction step, we retrieved 34 genomic bins representing: 27 HQ MAGs regarding MIMAG criteria [3], which are > 90% complete and < 5% contaminated, as well as they present ribosomal genes and at least 18 canonical tRNAs; and seven MQ MAGs, which are > 50% complete and < 10% contaminated (Table 1, Figure 1). The frameshift correction step [14] applied to the initial genomic bins reduced insertion and deletion errors –the most common error in Nanopore sequencing– of the CanMAGs (Additional File 2). After this extra correction step, the completeness was either increased or maintained, transforming five MQ MAGs to HQ MAGs.

**Table 1.** Comparison of CanMAGs to their representatives in databases considering MIMAG criteria. HQ MAGs must have > 90% completeness, < 5% contamination, ribosomal rRNAs and at least 18 canonical tRNAs, regarding MIMAG criteria [REF]. All the CanMAGs are compared to their GTDB representative (Rep.), in exception of those MAGs considered novel species (taxonomy at the genus level: g_). For these, we used high completeness MAGs with > 95% ANI from the animal gut metagenome or UHGG as a reference. *Phocaeicola* species, classified as Bacteroides in NCBI [31] *Genome assemblies derived from pure culture **Genome assemblies with “complete” level in NCBI, so no gaps and with no unplaced scaffolds.

|  |  | Genome size<br>(Mbp) |  | % Completeness |  | % Contamination |  | Total rRNAs |  | Canonical tRNAs |  | N° of contigs |  |
| --- | --- | --- | --- | --- | --- | --- | --- | --- | --- | --- | --- | --- | --- |
| Taxonomy (GTDB) | CanMAG vs. Rep. Genome | Can<br>MAG | Rep. | Can<br>MAG | Rep. | Can<br>MAG | Rep. | Can<br>MAG | Rep. | Can<br>MAG | Rep. | Can<br>MAG | Rep. |
| CanMAG vs. Short-read MAG Rep. |  |  |  |  |  |  |  |  |  |  |  |  |  |
| *Phascolarctobacterium_A* sp900544885 | CanMAG_01 vs GCA_900544885.1 | 2.09 | 1.75 | 99.85 | 98.65 | 2 | 1.5 | 15 | 1 | 20 | 18 | 1 | 87 |
| *Clostridium_Q* sp000435655 | CanMAG_02 vs GCA_000435655.1 | 3.11 | 2.73 | 94.79 | 96.68 | 0 | 0 | 15 | 0 | 20 | 14 | 12 | 149 |
| *g__Erysipelatoclostridium; s__* | CanMAG_07 vs Bissell_001 | 2.13 | 1.77 | 90.54 | 90.25 | 0 | 0 | 15 | 1 | 20 | 6 | 28 | 418 |
| *Blautia* sp000432195 | CanMAG_10 vs GCA_000432195.1 | 3.08 | 3.02 | 93.39 | 97.52 | 2.55 | 1.27 | 9 | 0 | 19 | 17 | 45 | 70 |
| *Blautia* sp900556555 | CanMAG_12 vs Peterbilt_039 | 2.96 | 2.65 | 97.64 | 97.5 | 0 | 0.48 | 12 | 0 | 20 | 14 | 15 | 106 |
| *Blautia_A* sp900541345 | CanMAG_14 vs GCA_900541345.1 | 2.73 | 2.69 | 97.97 | 95.85 | 0 | 0 | 18 | 0 | 20 | 16 | 10 | 160 |
| *g__Schaedlerella; s__* | CanMAG_19 vs Scrappy_009 | 2.47 | 2.32 | 94.63 | 95.35 | 0 | 0.58 | 15 | 0 | 20 | 15 | 5 | 51 |
| *g__UMGS966; s__* | CanMAG_21 vs Oklahoma_026 | 1.97 | 1.45 | 94.3 | 84.73 | 0 | 0.06 | 12 | 2 | 20 | 14 | 5 | 223 |
| *Phocaeicola* sp900546645 | CanMAG_25 vs GCA_900546645.1 | 3.40 | 2.82 | 98.32 | 92.56 | 0.82 | 0.87 | 18 | 1 | 20 | 15 | 33 | 131 |
| *Phocaeicola* sp900556845 | CanMAG_26 vs Flurry_018 | 3.31 | 2.55 | 98.74 | 96.81 | 0.52 | 0.15 | 24 | 0 | 20 | 13 | 3 | 167 |
| *Prevotellamassilia* sp000437675 | CanMAG_27 vs GCA_000437675.1 | 3.38 | 2.62 | 98.02 | 97.55 | 2.23 | 0.37 | 24 | 0 | 20 | 16 | 3 | 172 |
| *Prevotellamassilia* sp900541335 | CanMAG_28 vs GCA_900541335.1 | 2.72 | 2.42 | 97.65 | 96.13 | 0 | 0.05 | 21 | 0 | 20 | 16 | 1 | 95 |
| *g__Sutterella; s__* | CanMAG_31 vs MGYG-HGUT-01574 | 2.89 | 1.14 | 96.21 | 78.72 | 1.24 | 0.31 | 28 | 0 | 20 | 14 | 2 | 24 |
| *g__Succinivibrio; s__* | CanMAG_32 vs Freddie_038 | 2.04 | 1.74 | 98.68 | 97.5 | 0 | 0 | 22 | 0 | 20 | 14 | 1 | 185 |
| *Fusobacterium_B* sp900554885 | CanMAG_34 vs Glacier_008 | 2.06 | 1.57 | 96.63 | 100 | 1.28 | 0 | 21 | 1 | 20 | 19 | 47 | 220 |
| *g__Holdemanella; s__* | mq CanMAG_08 vs Bissell_031 | 2.40 | 1.74 | 85.25 | 97.71 | 1.99 | 1.65 | 20 | 0 | 18 | 12 | 21 | 179 |
| *UBA9502* sp900538475 | mq CanMAG_18 vs GCA_900538475.1 | 2.58 | 3.04 | 73.43 | 99.37 | 0.95 | 0 | 18 | 0 | 20 | 18 | 9 | 46 |
| *Faecalibacterium* sp900540455 | mq CanMAG_20 vs GCA_900540455.1 | 2.38 | 2.3 | 82.09 | 99.81 | 1.02 | 0.68 | 18 | 2 | 20 | 18 | 24 | 71 |
| *g__Bacteroides; s__* | mq CanMAG_29 vs Scooby_030 | 2.22 | 2.57 | 75.11 | 97.01 | 0 | 0.45 | 18 | 1 | 19 | 16 | 1 | 186 |
| **CanMAG vs. Short-read WGS Rep.** |  |  |  |  |  |  |  |  |  |  |  |  |  |
| *Catenibacterium* sp000437715 | CanMAG_05 vs GCF_004168205.1* | 2.57 | 2.54 | 98.46 | 100 | 0 | 0 | 30 | 10 | 20 | 20 | 2 | 212 |
| *Allobaculum stercoricanis* | CanMAG_06 vs GCF_000384195.1* | 2.18 | 2.05 | 97.64 | 98.11 | 2.29 | 2.29 | 15 | 21 | 20 | 20 | 33 | 3 |
| *Enterocloster* sp001517625 | CanMAG_15 vs GCF_001517625.2* | 3.64 | 3.41 | 98.92 | 99.37 | 1.67 | 0 | 15 | 8 | 20 | 20 | 62 | 7 |
| *Faecalimonas umbilicata* | CanMAG_16 vs GCF_004346095.1* | 2.42 | 3.08 | 91.8 | 99.37 | 0 | 0 | 18 | 2 | 20 | 18 | 20 | 65 |
| *Ruminococcus_B gnavus* | CanMAG_17 vs GCF_002959615.1* | 3.52 | 3.57 | 94.63 | 99.42 | 3.63 | 0 | 15 | 10 | 19 | 20 | 60 | 7 |
| *Megamonas funiformis* | CanMAG_22 vs GCF_000245775.1* | 2.38 | 2.56 | 94.94 | 99.68 | 3.16 | 0.63 | 9 | 12 | 19 | 18 | 51 | 13 |
| *Phocaeicola coprocola* | CanMAG_23 vs GCF_000154845.1* | 3.61 | 4.3 | 96.64 | 98.88 | 0.57 | 0 | 18 | 17 | 20 | 20 | 7 | 90 |
| *Phocaeicola plebeius* | CanMAG_24 vs GCF_000187895.1* | 3.31 | 4.42 | 97.57 | 99.25 | 0.56 | 0.68 | 18 | 14 | 20 | 19 | 2 | 19 |
| *Collinsella intestinalis* | CanMAG_33 vs GCF_000156175.1* | 2.18 | 1.81 | 99.19 | 99.19 | 0 | 0 | 12 | 11 | 20 | 20 | 49 | 3 |
| *Clostridium_U hiranonis* | mq CanMAG_03 vs GCF_000156055.1* | 2.62 | 2.48 | 91.38 | 100 | 6.29 | 0 | 27 | 3 | 10 | 14 | 68 | 26 |
| *Blautia_A* sp000433815 | mq CanMAG_13 vs GCF_005844445.1* | 2.68 | 3.48 | 65.14 | 99.37 | 0 | 0 | 9 | 3 | 18 | 18 | 15 | 102 |
| *Sutterella wadsworthensis_A* | mq CanMAG_30 vs GCF_000297775.1* | 3.27 | 2.73 | 98.28 | 98.14 | 6.58 | 0.62 | 17 | 14 | 20 | 20 | 68 | 11 |
| **CanMAG vs. Complete genome Rep.** |  |  |  |  |  |  |  |  |  |  |  |  |  |
| *Enterococcus_B hirae* | CanMAG_04 vs GCF_000271405.2** | 2.81 | 2.83 | 99.13 | 99.63 | 0 | 0 | 18 | 18 | 20 | 20 | 2 | 2 |
| *Blautia hansenii* | CanMAG_09 vs GCF_002222595.2** | 3.68 | 3.07 | 96.02 | 99.36 | 4.35 | 0 | 12 | 15 | 20 | 20 | 61 | 1 |
| *Blautia* sp003287895 | CanMAG_11 vs GCF_003287895.1** | 3.12 | 3.3 | 99.36 | 97.64 | 0 | 0.32 | 15 | 14 | 20 | 20 | 3 | 1 |

**Figure 1.**
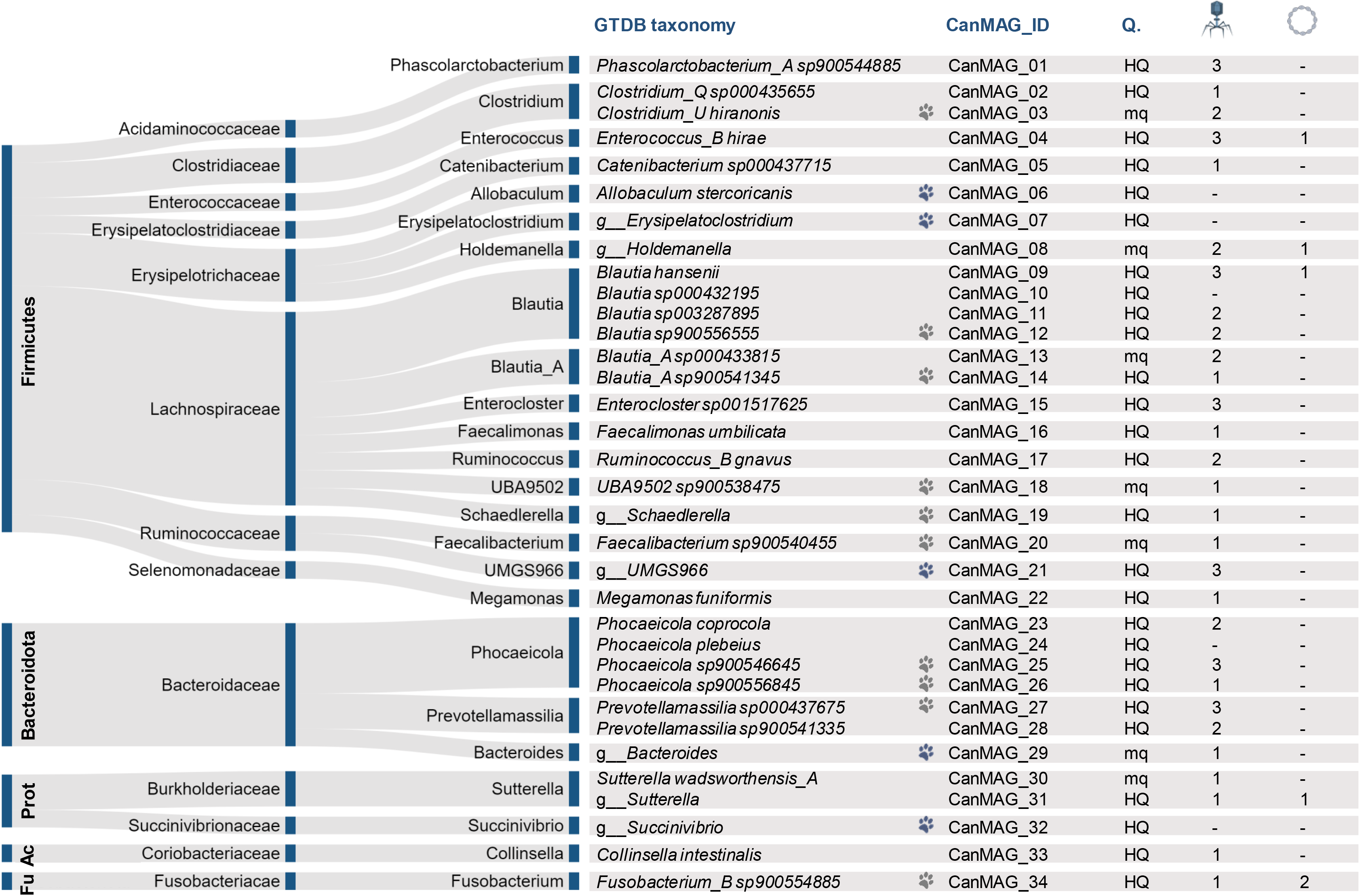
HQ and MQ CanMAGs from a canine fecal sample and their associated bacteriophages and plasmids. Reads with a taxonomy of “g_” are considered novel species by GTDB-tk. Blue paw indicates that the bacterial species has only been observed in dogs when assessing animal fecal microbiome; grey paw indicates that the bacterial species is more prevalent in dogs. HQ, high-quality MAG and mq, medium-quality MAG regarding MIMAG criteria [3]. All the predicted bacteriophages were integrated within the bacterial host chromosome. Fu.: Fusobacteriota, Ac.: Actinobacteriota, and Prot.: Proteobacteriota.

Most of the recovered CanMAGs belonged to Firmicutes phylum (n=21), followed by Bacteroidota (n=8) and Proteobacteriota (n=3). Overall, the most abundant genera recovered were: four *Blautia*, two *Blautia_A,* and two *Clostridium* species (Firmicutes); four *Phocaeicola* (former *Bacteroides* species [31]), and two *Prevotellamassilia* (Bacteroidota); and two *Sutterella* (Proteobacteriota) (Table 1, Figure 1). Even though eight of the CanMAGs were considered novel species by GTDB-tk, CanMAG bacterial species had been previously detected in other metagenome collections, and five exclusively in canine feces (Figure 1, Additional File 3).

The representative genomes in public databases for CanMAG bacterial species were (Table 1): i) short-read MAGs (n=19; 10 from fecal catalogs and 9 from GTDB); ii) genome assemblies from pure cultures (contig- or scaffold-level assemblies; n=12); or complete genomes (n=3).

Short-read MAGs representative genomes presented contig- or scaffold- level assemblies (24 to 223 contigs, mean=144) and had from 0 to 2 ribosomal genes and from 6 to 19 canonical tRNAs (mean=15). When compared to them, HQ CanMAGs recovered more ribosomal genes and canonical tRNA genes. Moreover, they presented a more contiguous assembly with larger genome sizes.

Genomes assemblies from pure cultures were also contig- or scaffold- level (3 to 212 contigs, mean=46), had from 2 to 21 ribosomal genes (mean=10), and from 18 to 20 canonical tRNAs (mean=19). When compared to them, HQ CanMAGs usually recovered more ribosomal genes (7 out of 9 bacterial species), even though in some cases CanMAG genome assembly was less contiguous. Only for *Allobaculum stercoricanis* and *Megamonas funiformis*, the representative genome derived from type strain material (GCF_000384195.1 and GCF_000245775.1, respectively) harbored more ribosomal genes than the CanMAGs.

Complete genome assemblies (reference genomes) were single-contig, presented all the ribosomal genes and the 20 canonical tRNAs. For *Enterococcus hirae*, we identified the same number of ribosomal genes as the reference genome (GCF_000271405.2).

Besides linking long contigs to retrieve HQ MAGs, Hi-C proximity ligation linked some potential plasmids to their bacterial host (Figure 1). We identified six potential plasmids linked to *Enterococcus hirae*, *g_ Holdemanella*, *Blautia hansenii* CanMAGs, *g_ Sutterella*, and two plasmids to *Fusobacterium_B sp900554885* CanMAG (Figure 1, Additional File 4). They presented an increased coverage compared to their bacterial host chromosome, and five of them were circular. Moreover, the plasmids contained typical plasmid or mobilome associated genes and blasted to previously identified plasmids –despite usually with a low coverage–. Moreover, one of the plasmids (PL2-CanMAG_34 in *Fusobacterium_B sp900554885*) harbored an antimicrobial resistance gene to Lincosamide (l*inA*).

### Linking prophages to their bacterial host on dog fecal microbiome

We detected 51 bacteriophages in the CanMAGs. The bacteriophages were integrated within the bacterial chromosome (prophages) rather than in free viral particles (Figure 1, Table 2): 30 were HQ (> 90% completeness), and 21 were genome-fragments with > 50% completeness (as defined by MIUViG criteria [29]) (Table 2). Low-quality predicted bacteriophages (as determined by Checkv) were not included in this analysis. We named these bacteriophages (BP), regarding their CanMAG bacterial host as follows BPX-CanMAG_XX.

**Table 2.** Predicted bacteriophages in CanMAGs: main characteristics and clustering information. Most of the predicted bacteriophages (BP) were integrated into the CanMAG bacterial genome and double-stranded DNA. We clustered them together with a Gut Phage database (GPD) subset to create viral clusters (VC). BP sequences were classified as: Clustered (C.), when confidently grouping in a VC; Outlier (Out.), when despite some links to a VC, the association was not statistically significant; Overlap (Ovl.), when the BP was linked to two or more VCs; or Singleton (S.), when it did not match any VC. % Compl. is % completeness, as assessed by Checkv. Details on the VCs can be found in Additional File 5. *GPD Bacterial host: predicted bacterial host for GPD representatives within a specific VC, if variable taxa, we state the lowest shared taxonomic information. N/D Not determined: no reported bacterial host in GPD.

| Bacterial host (here) | BP ID | VC | VC status | VC Size | BP length | % Compl. | gene count | viral genes | host genes | GPD Bacterial host* |
| --- | --- | --- | --- | --- | --- | --- | --- | --- | --- | --- |
| **Firmicutes** |  |  |  |  |  |  |  |  |  |  |
| *Enterocloster* sp001517625 | BP1-CanMAG_15 | VC_183 | C. | 11 | 25,334 | 65.49 | 35 | 14 | 0 | *Lachnospiraceae* |
| *UBA9502* sp900538475 | BP1-CanMAG_18 | VC_183 | C. | 11 | 39,523 | 100 | 66 | 16 | 0 | *Lachnospiraceae* |
| *Blautia* sp003287895 | BP1-CanMAG_11 | VC_301 | C. | 5 | 28,487 | 83.11 | 38 | 16 | 0 | *Lachnospiraceae* |
| *Blautia* sp900556555 | BP1-CanMAG_12 | VC_344 | C. | 9 | 34,086 | 90.85 | 55 | 19 | 0 | *Lachnospiraceae* |
| *Blautia* sp900556555 | BP2-CanMAG_12 | VC_344 | C. | 9 | 36,598 | 100 | 50 | 10 | 0 | *Lachnospiraceae* |
| *Blautia_A* sp000433815 | BP1-CanMAG_13 | VC_241 | C. | 7 | 26,155 | 74.49 | 53 | 11 | 0 | N/D |
| *Blautia hansenii* | BP1-CanMAG_09 | VC_342 | C./S. | - | 151,986 | 89.15 | 226 | 40 | 7 | *Blautia hansenii* |
| *g_UMGS966; s_* | BP1-CanMAG_21 | VC_267 | C. | 8 | 47,108 | 100 | 65 | 21 | 2 | *Ruminococcaceae* |
| *Clostridium_Q* sp000435655 | BP1-CanMAG_02 | VC_254 | C. | 5 | 53,237 | 100 | 74 | 15 | 3 | N/D |
| *Clostridium_U hiranonis* | BP1-CanMAG_03 | VC_347 | C. | 3 | 34,195 | 51.54 | 55 | 22 | 1 | *Clostridium_U hiranonis* |
| *Blautia_A* sp000433815 | BP2-CanMAG_13 | VC_553 | C. | 3 | 150,650 | 100 | 143 | 1 | 66 | N/D |
| *Megamonas funiformis* | BP1-CanMAG_22 | VC_253 | C. | 5 | 35,900 | 100 | 57 | 16 | 1 | *Megamonas funiformis* |
| *Catenibacterium* sp000437715 | BP1-CanMAG_05 | VC_217 | C. | 27 | 45,860 | 97.95 | 53 | 23 | 1 | *Firmicutes* |
| *g_Holdemanella; s_* | BP1-CanMAG_08 | VC_217 | C. | 27 | 44,640 | 88.4 | 60 | 21 | 3 | *Firmicutes* |
| *Enterocloster* sp001517625 | BP2-CanMAG_15 | VC_217 | C. | 27 | 27,920 | 59.55 | 33 | 19 | 2 | *Firmicutes* |
| *Phascolarctobacterium_A* sp900544885 | BP2-CanMAG_01 | VC_555 | C. | 4 | 39,056 | 95.43 | 59 | 22 | 0 | *Negativicutes* |
| *Phascolarctobacterium_A* sp900544885 | BP1-CanMAG_01 | VC_036 | C. | 4 | 57,434 | 100 | 92 | 44 | 0 | *Negativicutes* |
| *Faecalibacterium* sp900540455 | BP1-CanMAG_20 | VC_403 | C. | 16 | 34,244 | 97.95 | 53 | 22 | 1 | N/D |
| *Blautia hansenii* | BP2-CanMAG_09 | VC_552 | C. | 3 | 3,767 | 90.52 | 5 | 1 | 0 | *Lachnospiraceae* |
| *Ruminococcus_B gnavus* | BP1-CanMAG_17 | VC_552 | C. | 3 | 6,213 | 100 | 10 | 2 | 0 | *Lachnospiraceae* |
| *Enterocloster sp001517625* | BP3-CanMAG_15 | VC_554 | C. | 7 | 191,453 | 68.4 | 258 | 1 | 13 | N/D |
| *Blautia hansenii* | BP3-CanMAG_09 | - | Out. | - | 25,525 | 51.57 | 20 | 1 | 3 | - |
| *Blautia sp003287895* | BP2-CanMAG_11 | - | Out. | - | 19,133 | 100 | 17 | 7 | 0 | - |
| *Blautia_A sp900541345* | BP1-CanMAG_14 | - | Out. | - | 27,724 | 58.84 | 40 | 12 | 0 | - |
| *g_UMGS966; s_* | BP2-CanMAG_21 | - | Out. | - | 40,694 | 89.88 | 61 | 27 | 1 | - |
| *g_UMGS966; s_* | BP3-CanMAG_21 | - | Out. | - | 29,305 | 66.4 | 45 | 12 | 1 | - |
| *Clostridium_U hiranonis* | BP2-CanMAG_03 | - | Ovl. | - | 41,047 | 75.27 | 70 | 23 | 1 | - |
| *Phascolarctobacterium_A sp900544885* | BP3-CanMAG_01 | - | Ovl. | - | 22,711 | 54.16 | 34 | 13 | 1 | - |
| *Ruminococcus_B gnavus* | BP2-CanMAG_17 | - | Ovl. | - | 36,619 | 95.92 | 67 | 20 | 0 | - |
| *Enterococcus_B hiraе* | BP1-CanMAG_04 | - | S. | - | 32,704 | 50.36 | 38 | 8 | 3 | - |
| *Enterococcus_B hiraе* | BP2-CanMAG_04 | - | Ovl. | - | 41,858 | 100 | 58 | 34 | 0 | - |
| *Enterococcus_B hiraе* | BP3-CanMAG_04 | - | Out. | - | 34,545 | 90.55 | 50 | 9 | 3 | - |
| *Faecalimonas umbilicata* | BP1-CanMAG_16 | - | Ovl. | - | 33,688 | 83.74 | 57 | 24 | 0 | - |
| *g_Schaedlerella; s_* | BP1-CanMAG_19 | - | Out. | - | 37,489 | 93.41 | 51 | 13 | 1 | - |
| *g_Holdemanella; s_* | BP2-CanMAG_08 | - | Ovl. | - | 33,282 | 96.2 | 62 | 21 | 0 | - |
| **Bacteroidota** |  |  |  |  |  |  |  |  |  |  |
| *Phocaeicola sp900546645* | BP1-CanMAG_25 | VC_219 | C. | 12 | 34,229 | 92.18 | 44 | 18 | 2 | *Phocaeicola* |
| *Phocaeicola sp900556845* | BP1-CanMAG_26 | VC_318 | C. | 10 | 47,132 | 100 | 63 | 9 | 1 | *Phocaeicola* |
| *Phocaeicola coprocola* | BP2-CanMAG_23 | VC_544 | C. | 4 | 57,738 | 100 | 63 | 4 | 3 | *Bacteroidaceae* |
| *Phocaeicola sp900546645* | BP2-CanMAG_25 | VC_544 | C. | 4 | 44,212 | 98.3 | 54 | 7 | 1 | *Bacteroidaceae* |
| *Phocaeicola sp900546645* | BP3-CanMAG_25 | VC_545 | C. | 18 | 58,284 | 91.08 | 55 | 9 | 2 | *Phocaeicola* |
| *Prevotellamassilia sp000437675* | BP3-CanMAG_27 | VC_547 | C. | 3 | 31,919 | 75.45 | 43 | 2 | 1 | *Bacteroidaceae* |
| *Phocaeicola coprocola* | BP1-CanMAG_23 | VC_508 | C. | 11 | 59,043 | 100 | 74 | 10 | 3 | *Bacteroidales* |
| *Prevotellamassilia sp000437675* | BP2-CanMAG_27 | VC_510 | C. | 13 | 44,671 | 92.29 | 47 | 5 | 5 | *Bacteroidaceae* |
| *Prevotellamassilia* sp000437675 | BP1-CanMAG_27 | VC_405 | C. | 5 | 37,057 | 74.49 | 49 | 8 | 0 | N/D |
| *g__Bacteroides; s__* | BP1-CanMAG_29 | VC_488 | C. | 3 | 6,365 | 100 | 9 | 3 | 0 | N/D |
| *Prevotellamassilia* sp900541335 | BP1-CanMAG_28 | - | Out. | - | 20,636 | 54.51 | 16 | 1 | 4 | - |
| *Prevotellamassilia* sp900541335 | BP2-CanMAG_28 | - | Out. | - | 37,022 | 57.64 | 18 | 2 | 2 | - |
| **Fusobacteriota** |  |  |  |  |  |  |  |  |  |  |
| *Fusobacterium_B* sp900554885 | BP1-CanMAG_34 | VC_348 | C. | 12 | 43,899 | 100 | 75 | 13 | 2 | *Fusobacterium* |
| **Proteobacteriota** |  |  |  |  |  |  |  |  |  |  |
| *g__Sutterella; s__* | BP1-CanMAG_31 | VC_257 | C. | 7 | 42,692 | 90.3 | 72 | 27 | 0 | N/D |
| *Sutterella wadsworthensis_A* | BP1-CanMAG_30 | - | Ovl. | - | 45,521 | 95.47 | 65 | 27 | 2 | - |
| **Actinobacteriota** |  |  |  |  |  |  |  |  |  |  |
| *Collinsella intestinalis* | BP1-CanMAG_33 | - | S. | - | 2,515 | 60.44 | 2 | 1 | 0 | - |

When clustering our bacteriophages together with a subset of the Gut Phage Database (GPD, [44]) –containing 682 bacteriophage sequences–, we obtained 27 viral clusters (VC) (Table 2, Figure 2, Additional File 5). Viral clusters grouped bacteriophages with similar genome sizes and bacterial hosts. Bacteriophage genome sizes ranged from 2,515 to 191,453 bp (Figure 2A). Thirty-three bacteriophages were distributed and clustered in 27 viral clusters, containing three to 27 bacteriophage sequences (Figure 2B, Additional File 5). The remaining 18 bacteriophages were classified as: outliers (n=9), when they were attached to a VC, but not statistically significant; overlap (n=7), when they presented overlapping genes between two or more VC; and singletons (n=2), when they did not cluster with anything else.

**Figure 2.**
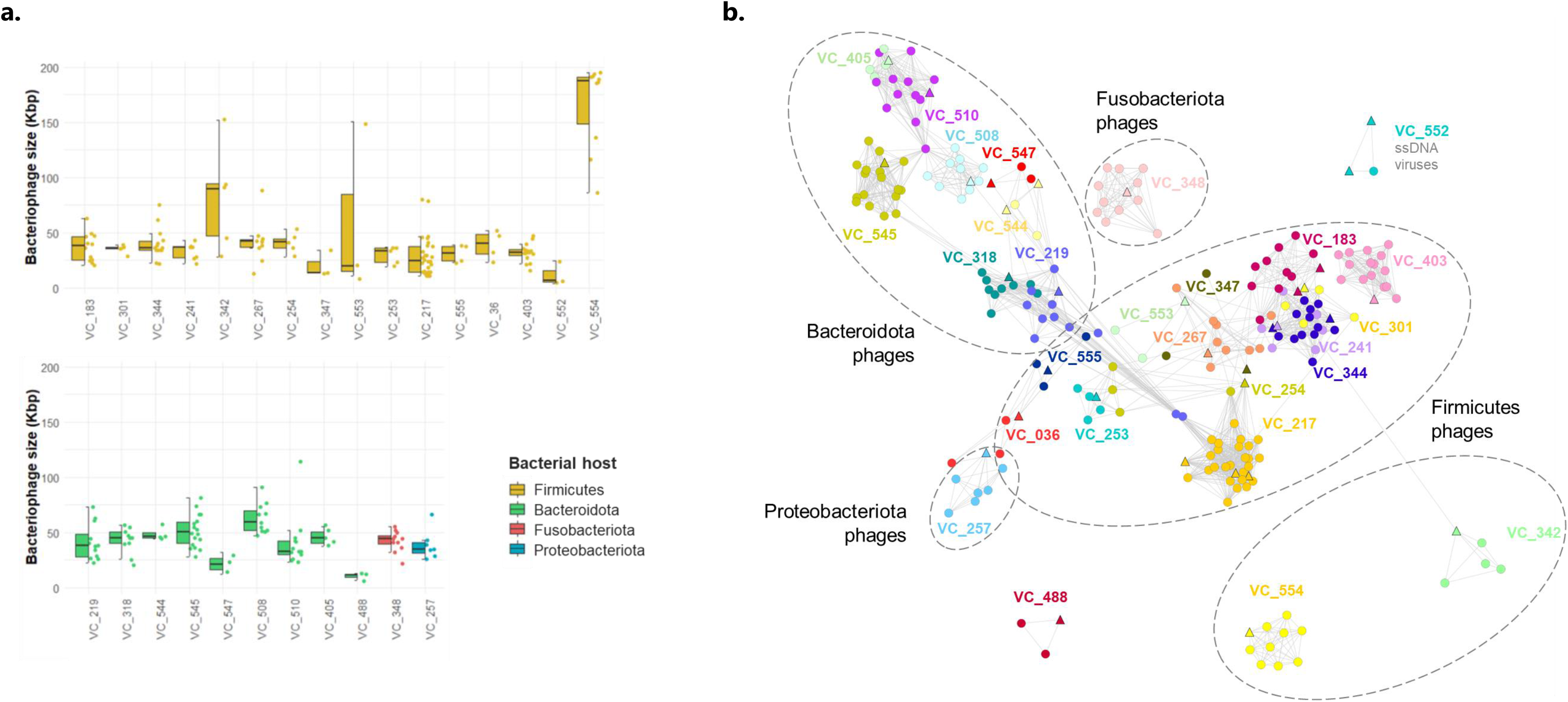
Analysis of the 27 viral clusters (VC) that included our CanMAG bacteriophages. Both figures contain data from the 33 clustered CanMAG bacteriophages and the representatives from GPD grouping together within the same VC. A) Boxplots representing the bacteriophages genome sizes within the cluster colored by bacterial host phylum. B) Viral clusters network. For visualization purposes, the predicted bacteriophages from CanMAGs are pictured as triangles, and the bacteriophages from the Gut Phage database, as circles.

Our results provided novel bacterial host information for eight out of the 27 VC including GPD viral genomes (N/D in GPD Bacterial host in Table 2): VC_241, VC_254, VC_553, VC_403, VC_554, VC_405, VC_488, and VC_257. Three viral clusters shared a specific bacterial host: VC_253 contained bacteriophages only observed in *Megamonas funiformis*; VC_342, in *Blautia hansenii*; and VC_347, in *Clostridium hiranonis*. Four viral clusters shared the same bacterial host at the genus level: VC_219, VC_545, and VC_318 contained bacteriophages only observed in *Phocaeicola* genus; and VC_348, in *Fusobacterium*. The remaining viral clusters grouped bacteriophages with a broader range of bacterial hosts (family or above).

Finally, all the bacteriophages were predicted to be integrated, except BP3-CanMAG_15 that was circular, lytic, and clustered together with other GPD bacteriophages in VC_554 despite harboring only one viral protein, probably representing another extra-chromosomal element rather than a lytic virus. Besides, most of the predicted prophages were double-stranded DNA, except three that Virsorter2 predicted as single-stranded DNA: BP1-CanMAG_17 (*Ruminococcus_B gnavus*) and BP2-CanMAG_09 (*Blautia hansenii*), which were clustering together in VC552; and BP1-CanMAG_33 (*Collinsella intestinalis*), which was a singleton.

### CanMAGs recovered more mobilome-associated gene functions when compared to short-read MAGs representatives

We were interested in assessing long-read metagenomics to recover overall mobilome-associated gene functions, typically encoded by mobile genetic elements with repetitive regions that are difficult to characterize due to collapse of short reads.

Seventeen out of 27 HQ CanMAGs recovered more mobilome-associated gene functions (Mobilome COG category) when compared to their bacterial representatives. Three HQ CanMAGs recovered the same MGEs as their genome representatives (Additional File 6). They represented *Allobaculum stercoricanis, Blautia_A sp900541345, g_UMGS966* that were species predicted to be more prevalent in the canine fecal environment (Figure 1).

We used the *Phocaeicola* genus (includes some former *Bacteroides* species [31]) as an example to further assess mobilome functions since we identified four different bacterial species within this dog fecal metagenome, and they presented abundant mobilome-related functions. Thus, we computed the pangenome for the *Phocaeicola* genus, including the CanMAGs, the GTDB reference genomes, and a MAG from the UHGG and the dog gut catalog per bacterial species (when available).

*Phocaeicola* CanMAGs presented more mobilome-associated functions when compared to short-read MAGs, being the proportion more similar to that previously reported on genome assemblies from type strain material (pure culture) rather than that found in short-read MAGs (complex microbial community) (Figure 3A).

**Figure 3.**
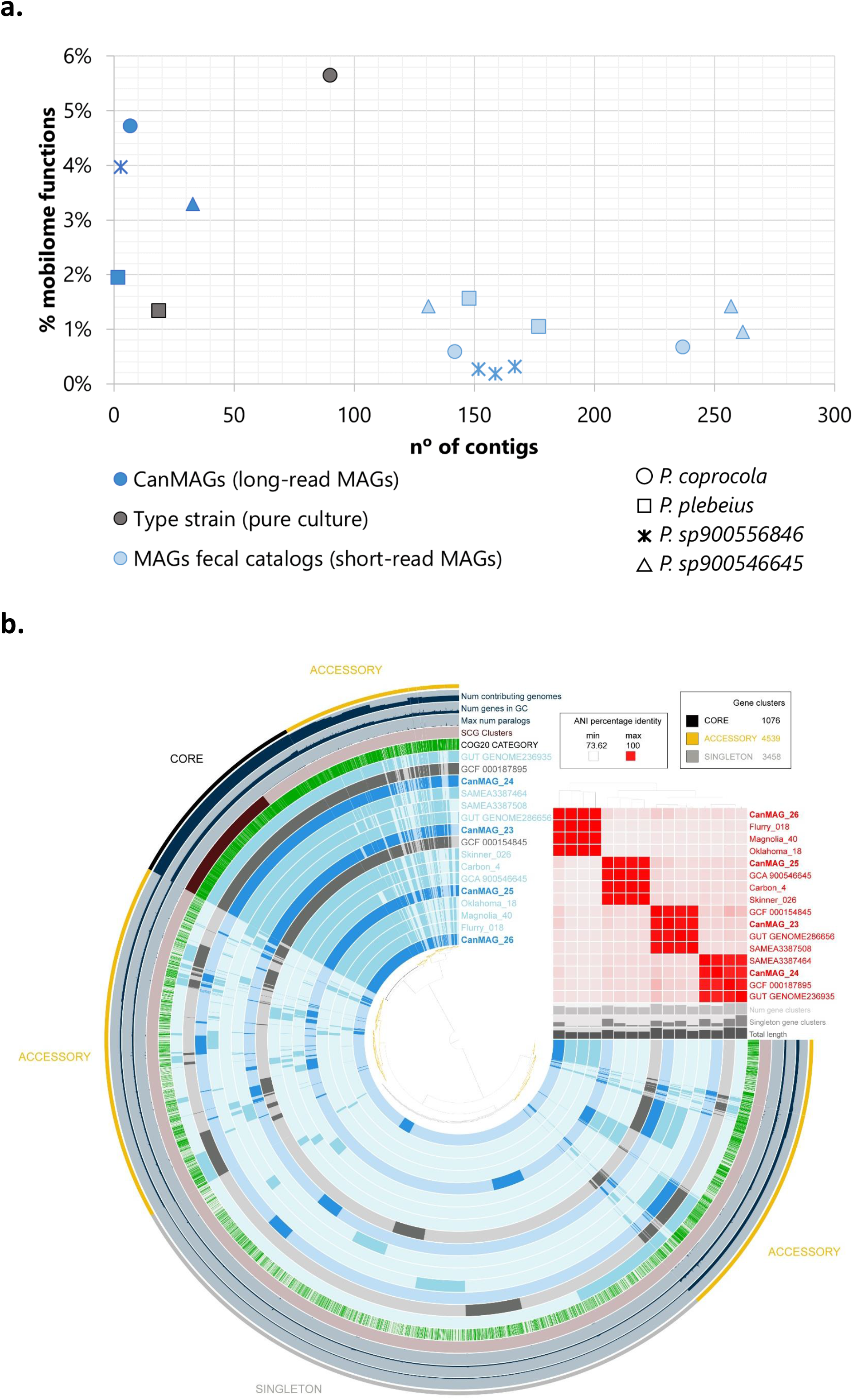
*Phocaeicola* species comparison to published bacterial genomes. We included the four *Phocaeicola* species CanMAGs, their respective GTDB reference genome, and a MAG from the UHGG and the dog gut catalog per bacterial species (when available). A) Percentage of mobilome functions per genome considering the contiguity of the assembly. Colors indicate the source., whereas forms indicate the bacterial species. B) Pangenome visualization for Phocaeicola species. ANI coloring value lower limit is genus-level threshold [52] CanMAGs are colored with a darker blue, complete genomes with grey, and short-read MAGs, lighter blue.

Apart from the mobilome, CanMAGs presented similar gene functions and gene clusters compared to their representatives as observed in the accessory genome pattern in the pangenome visualization (Figure 3B). Moreover, each CanMAG clustered together with its bacterial species representatives with ANIs > 95% (species threshold [26]).

## Discussion

The metagenomics field can benefit from long-read sequencing technologies since they span long DNA stretches and repetitive regions to retrieve complete ribosomal genes and mobile genetic elements (MGEs) with genome context to facilitate highly contiguous assemblies. Hi-C proximity ligation can help disentangle a complex microbiome by capturing *in vivo* interactions that can be used to bin genome contigs or associate extra-chromosomal elements to their bacterial host [15,16]. In this study, we have combined both approaches retrieving 27 HQ MAGs and seven MQ MAGs from the fecal microbiome of a healthy dog.

We previously used the same long-read metagenomics data to recover eight HQ MAGs as single-contigs by combining assembly outputs from different datasets (all data, 75% data, HMW data) [19]. Here, we further improved the contiguity of the previous metagenome assembly (using “all data” dataset) by binning the long-contigs with Hi-C proximity ligation data. We retrieved different bacterial species within the same genera, as seen for *Phocaeicola* and *Blautia* species. These results agree with the recently described species-level MAGs from a sheep fecal microbiome combining PacBio HiFi reads and Hi-C metagenomics binning [18]. Another potential binning approach is using bioinformatics software, but the most common binning software were developed for short-read metagenomics. Further steps will evaluate the performance of recently developed long-read binning software [45].

All the 27 HQ MAGs identified fulfilled the MIMAG criteria being > 90% complete and < 5% contaminated and presenting ribosomal genes and at least 18 canonical tRNAs [3]. None of the previously reported canine MAGs –for the bacterial species identified here-fulfilled MIMAG criteria for HQ MAG, despite being highly complete [28,29]. In fact, short-read MAGs usually lack most of the ribosomal genes, which end up collapsed and unassembled since they are repeated and highly conserved [5]. In the Unified Human Gastrointestinal Genomes (UHGG) catalog, they found 3,207 species-level non-redundant genomes that were highly complete (> 90% completeness), but only 38 of them were MAGs that met the HQ criteria regarding MIMAG [2]. Therefore, our approach allows to retrieve not only highly complete MAGs, but also HQ MAGs considering MIMAG criteria.

Apart from the correct assembly and location of the ribosomal genes, long-read metagenomics can assemble and locate MGEs. Compared to short-read MAGs, the long-reads CanMAGs recovered larger percentages of mobilome COG gene functions (similar to genomes recovered from pure cultures of type material). This fact agrees with previous long-read metagenomics surveys that succeeded in assembling highly repetitive bacterial genomes [8,9,46] –MGEs commonly contain many repetitions– whereas short-read assemblies commonly break at that point [9]. Even considering that differences in mobilome functions can be due to the mobile nature of their genes (e.g., mobilization due to a horizontal gene transfer event), Prokka annotated the same MGEs in *Allobaculum stercoricanis* CanMAG as in the representative bacterial genome, which is an isolate from canine feces (GCF_000384195.1), a result that further validates our approach.

Long read metagenomics is likely to become a comprehensive approach to ascertain the bacterial host to bacteriophages. We clustered 51 prophages (with > 50% completeness) within the CanMAGs with a subset of the Gut Phage Database [44] and identified their host range. We found three species-specific bacteriophages for *Megamonas funiformis*, *Blautia hansenii*, and *Clostridium hiranonis*. Furthermore, we reported novel bacterial host information for the other eight bacteriophages. Overall, most of the prophages presented a broad spectrum of bacterial hosts, as suggested by previous researchers [47]. This fact contrasts with Gut Phage database findings, where most of the viral clusters were predicted to be species-specific, although > 70% of bacteriophages lacked host information [44].

Apart from the experimental binning of the genomes, Hi-C proximity ligation cross-link extra-chromosomal elements within a single cell [16,17,48–50]. We linked six potential plasmids to their bacterial host. We suspect that we may have missed some plasmids due to the use of the Ligation sequencing kit rather than the Rapid sequencing kit for the Nanopore library preparation, as it has recently been reported [51], after we had our experimental data. Since we aimed to retrieve longer reads, the Ligation sequencing kit was our first choice rather than the Rapid sequencing kit, which produces shorter reads because it uses transposase fragmentation to insert the adapters. If aiming to assess links between extra-chromosomal elements and their hosts, we would also recommend evaluating the use of the Rapid sequencing kit despite the shorter read length, which should be compensated with Hi-C binning data.

In conclusion, long-read metagenomics retrieved long contigs harboring complete assembled ribosomal operons, prophages, and other MGEs. Hi-C binned together the long contigs into HQ and MQ MAGs, some of them representing closely related species. Moreover, it also linked plasmids to their bacterial host. Long-read metagenomics and Hi-C binning are likely to become a comprehensive approach to discovering HQ MAGs and assigning extra-chromosomal elements to the bacterial host.

## Declarations

### Ethics approval and consent to participate

Not applicable

### Consent for publication

Not applicable

### Availability of data and materials

An overview of the scripts used to analyze the data is at Additional File 1. The final CanMAGs are available on Zenodo: 10.5281/zenodo.5055248. The raw fast5 files and the HQ MAGs are available on ENA under Bioproject PRJEB42270 (submission in process).

### Authors’ contributions

OF and AC conceptualized the study and designed the experiment. DP extracted the DNA, performed the sequencing libraries, the nanopore sequencing and the Hi-C proximity ligation protocol. AC performed the metagenome assembly and correction. AC analyzed and interpreted the data. JV and NF analyzed the plasmid data. NF performed the pangenome analysis. AC wrote the main manuscript text. OF, NF, DP and JV substantially revised the work. All the authors have approved the submitted version.

## Acknowledgments

We would like to thank Justa Martín, from Vetgenomics for the initial support on the Hi-C procedure. We would like to thank also Ivan Liahcko and Gherman Uritskiy from Phase Genomics for their support on the Hi-C data analysis.

## Additional Information

**Additional File 1. Bioinformatics workflow overview.** The file contains information on the software used and their versions, as well as the commands and the specific options to perform the bioinformatics analysis used here.

**Additional File 2. Frameshift-correction of CanMAGs.** Several quality values associated to MAG quality were assessed before and after the frameshift correction step. Completeness (Comp.), Contamination (Cont.), and number of predicted genes (n° pred. Genes) are values from CheckM. MAG quality regarding MIMAG criteria.

**Additional File 3. CanMAG bacterial species prevalence in animal and human gut catalogs.** The prevalence of bacterial species was compared to two fecal microbiome catalogs: animal gut metagenome (n=5,596) and UHGG catalogs (n=204,938). GTDB representative states the source of the representative genome on GTDB. Dog prevalence when comparing both catalogs.

**Additional File 4. Predicted plasmids linked to CanMAGs.** Predicted plasmids were manually checked assessing coverage (Cov.), circularity, predicted genes, and BLAST results.

**Additional File 5. Viral cluster network information.** Viral clusters included BPX-CanMAG_XX and a subset of Gut Phage Database viral genomes.

**Additional File 6. Mobilome functions in CanMAGs *vs*. representative genome.** Mobilome functions and gene products annotated by Prokka against COGs database. In green, CanMAGs with more mobilome functions. *Representative genomes derived from pure culture and **Representative genomes with a “complete genome” assembly level in NCBI. The remaining representative genomes are short-read MAGs from environmental sources

