## Additional File 1 for "Novel canine high-quality metagenome-assembled genomes, prophages, and host-associated plasmids by long-read metagenomics together with Hi-C proximity ligation"

### Additional File 1. Bioinformatics workflow overview

Anna Cusco

July 2021

This workflow aims to provide an overview of the analytical workflow used in "Novel canine high-quality metagenome-assembled genomes, prophages, and host-associated plasmids by long-read metagenomics together with Hi-C proximity ligation".

We have analyzed a **fecal microbiome** sample from a healthy dog. We have extracted the DNA from this sample using two kits from Zymobiomics:

- **Quick-DNA HMW MagBead** for High-molecular Weight DNA (without bead-beating).
- **DNA Miniprep Kit** for standard microbiome DNA extraction (with bead-beating)

Each of the DNA extractions has been run in an independent R9.4.1 flowcell in a MinION using SQK-LSK109 sequencing kit. We have merged the data and worked with the whole dataset.

Please consider checking Additional File 1 of doi.org/10.1186/s12864-021-07607-0, for the workflow of the 1) read pre-processing and 2) metagenome assembly and polishing steps. Here we list all the software used, but we provide the details of the analytical workflow from CanMAG characterization.

1. Raw read pre-processing software:

- **Guppy** 3.4.5 to basecall fast5 reads with high accuracy basecalling mode (dna_r9.4.1_450bps_hac.cfg)
- **Porechop** 0.2.4 to trim the adapters
- **Canu** 2.0 to correct raw reads

2. Metagenomics assembly and polishing:

- **Flye** 2.7-b1585 to perform metagenome assembly
- **Medaka** 1.0.1 to polish and correct the Flye-assembly
- **Diamond** 0.9.32 and **MEGAN-LR** 6.19.1 to correct insertions and deletions errors in HQ MAGs.

3. CanMAGs characterization

- **CheckM** 1.1.1 to assess completeness and contamination of the CanMAGs
- **GTDB-tk** 1.3.0 with GTDB taxonomy release 95 to assess the novelty and the taxonomy of HQ MAGs.
- **Prokka** 1.13.4 to annotate the CanMAGs.
- **Abricate** 0.9.8 to detect antimicrobial resistance genes using **CARD database**
- **Plasflow** 1.1.0 to predict potential plasmids within the CanMAGs

4. Pangenome of *Phocaeicola* genus

- **Anvi'o** 7 to perform the functional annotation (**COGs database**) and pangenomics analysis of *Phocaeicola* genus.

5. Bacteriophage characterization

- **VirSorter2** 2.1 and **Vibrant** 1.2.1 to predict potential bacteriophages within the CanMAGs
- **CheckV** 0.7.0 to assess the quality of the predicted bacteriophages, and trim host genomic regions.
- **vConTACT2** 0.9.19 to cluster the bacteriophages together with a subset of **Gut Phage Database** representatives.

#### 3) CanMAGs characterization

Combining a long-read metagenomics assembly with Hi-C proximity ligation data we retrieved 34 genomic bins representing: 27 HQ MAGs regarding and seven MQ MAGs. The genomic bins were retrieved by the Proximeta Cloud platform (Phase Genomics).

We assessed completeness and contamination with CheckM and predicted the taxonomy and potential novelty with GTDB-tk. We also annotated the genomes using Prokka.

```
## Run CheckM to assess completeness and contamination

checkm lineage_wf -t 8 -x fa CanMAGs/ /CheckM-out

## Run GTDB-tk to assess novelty and taxonomy

gtdbtk classify_wf --genome_dir CanMAGs/ \
--cpus 8 --out_dir /GTDB-tk-out

## Prokka to annotate the genome (CDS, ribosomal genes, tRNAs, etc.)

prokka --outdir prokka_CanMAG_XX --prefix CanMAG_XX --cpus 8 CanMAG_XX.fasta &

##PlasFlow to predict potential plasmids

PlasFlow.py --input CanMAG_XX.fasta --output CanMAG_XX-plasflow --threshold 0.7
```

### 4) Pangenome of *Phocaeicola* genus

We compared our Phocaeicola CanMAGs to previously reported MAGs from the same species: i) GTDB species representative genome, ii) the animal gut metagenome (Youngblut et al, 2020) and iii) the Unified Human Gastrointestinal Genome (UHGG) (Almeida et al, 2020).

We followed the pangenomics tutorial on Anvi'o website: https://merenlab.org/2016/11/08/pangenomics-v2/ (https://merenlab.org/2016/11/08/pangenomics-v2/)

We created an anvi'o contigs database for each one of the genome assemblies included in the pangenome (representative MAGs from public databases, and CanMAGs). This were the general commands followed:

```
##Simplify headers from each contig within fasta files

anvi-script-reformat-fasta <path>/.fasta -o <path>/.fasta -l 0 --simplify-names --report-file
<path>/.txt


##Transform fasta files in Anvi'o contigs DBs files

anvi-gen-contigs-database -f <path>/.fasta -n NAME -o <path>/.db


## Annotate functions using the COG20 database

anvi-run-ncbi-cogs -c <path>/.db -T 20 --temporary-dir-path <path>/xxx --search-with blastp

## Create an external genomes .txt file with names and path of each contig database within th
e pangenome

## Generate the genome storage file

anvi-gen-genomes-storage --external-genomes <path>/external-genomes.txt --gene-caller prodiga
l --output-file <path>/GENOMES.db

## Generate the pangenome database

anvi-pan-genome --genomes-storage <path>/GENOMES.db --use-ncbi-blast --minbit 0.5 --mcl-infla
tion 4 --project-name xxxxx --output-dir <path>/PAN.db --num-threads 20


## Compute ANI values

anvi-compute-genome-similarity --external-genomes <path>/external-genomes.txt --program pyANI
--output-dir <path>/pyANI --num-threads 6 --pan-db <path>/PAN.db

## Display the pangenome (Circular plot)

anvi-display-pan -p <path>/PAN.db -g <path>/GENOMES.db

## Core Genome Binning

anvi-get-sequences-for-gene-clusters -p <path>/PAN.db -g <path>/GENOMES.db -o <path>/core-gen
ome --min-num-genomes-gene-cluster-occurs x (x = total number of genomes analysed)

## Accessory Genome Binning

anvi-get-sequences-for-gene-clusters -p <path>/PAN.db -g <path>/GENOMES.db -o <path>/accessor
y-genome --max-num-genomes-gene-cluster-occurs y (y = total number of genomes analysed -1)

## Singleton Genome Binning

anvi-get-sequences-for-gene-clusters -p <path>/PAN.db -g <path>/GENOMES.db -o <path>/singleto
n-genome --max-num-genomes-gene-cluster-occurs 1 (maximum in 1 genome)

#Anvi'o Summary

anvi-summarize <path>/PAN.db -g <path>/GENOMES.db -C bins_NAME -o <path>/summary
```

### 5) Bacteriophage analysis

First step is the viral prediction in our final HQ and MQ CanMAGs. For doing so, we used both Virsorter2 and Vibrant.

```
##Install Virsorter2
conda create --prefix=virsorter-2 -c bioconda virsorter
source activate /scratch/114-canis.familiaris-MINION/condatools/virsorter-2

##Run Virsorter2 for each CanMAG
virsorter run -w CanMAG_XX.out -i CanMAG_XX.fa -j 4 --min-score 0.8 all

##Install Vibrant
conda create --prefix=Vibrant-1.2.1 -c bioconda vibrant
source activate /scratch/114-canis.familiaris-MINION/condatools/Vibrant-1.2.1

##Run Vibrant for each CanMAG
VIBRANT_run.py -i CanMAG_XX.fa -folder CanMAG_XX-vibrant
```

We concatenate each CanMAG predicted viruses file per software without merging the two output. So, we combine all the *final-viral-combined.fa* files for Virsorter2 in a single *_VS2* file and *XX-phages-combined.fa* files for Vibrant in a single *_VB* file.

We will proceed by running checkv, which performs a QC of the bacteriophages and trims the host parts present in the edges of prophages.

```
## Install checkv and set up the database
conda create --prefix=checkv -c conda-forge -c bioconda checkv
wget https://portal.nersc.gov/CheckV/checkv-db-v0.6.tar.gz
tar -zxvf checkv-db-v0.6.tar.gz
export CHECKVDB= /scratch/114-canis.familiaris-MINION/checkv_DB/checkv-db-v0.6 #path to where
you download the database

## Run checkv in the combined Virsorter2 file and in the Vibrant combined file.

checkv contamination final-viral-combined.fa output_checkv -t 16
checkv completeness final-viral-combined.fa output_checkv -t 16
checkv complete_genomes final-viral-combined.fa output_checkv
checkv quality_summary final-viral-combined.fa output_checkv
```

With checkv trimmed viruses and their associated quality measures, we create the final_predicted_viruses.fna file by:

- removing low-quality viruses
- when a virus was only predicted with one of the softwares, we include it
- when a virus was predicte with the two softwares, we keep the highest quality prediction (and the most complete).

Next step was clustering the CanMAG viruses, with publically available bacteriophages sequences. For this purpose we chose the “Gut Phage database” (GPD) (https://www.cell.com/cell/fulltext/S0092-8674(21)00072-6 (https://www.cell.com/cell/fulltext/S0092-8674(21)00072-6))

However, the Gut Phage database is too huge to perform the computationally-expensive step of viral protein clustering performed by vConTact-2. Subsequently, we first mapped at the nucleotide level our final_predicted_viruses.fna file against the complete Gut Phage database. Any GPD reference that was minimally mapping to our final_predicted_viruses.fna file was kept for the clustering step (682 bacteriophage sequences).

Finally, we combined final_predicted_viruses.fna file and the subset of GPD sequences to perform the viral clustering as detailed below.

```
##Minimap2 to map final_predicted_viruses.fna file against GPD_sequences.fa

minimap2 -cx map-ont GPD_sequences.fa final_predicted_viruses.fna file > final_predicted_viru
sesVSGPD.paf

##Create a list of the GPD IDs that mapped and subset the database

source activate /projects/114-canis.familiaris-MINION/condatools/seqkit-0.11/
seqkit grep -f GPD_hiting.txt GPD_sequences.fa -o GPD-hitting.fna
cat GPD-hitting.fna final_predicted_viruses.fna > final_predicted_viruses+GPD.fna

##Protein prediction of the viruses

source activate /scratch/114-canis.familiaris-MINION/condatools/prodigal-2.6.3/
prodigal -i final_predicted_viruses+GPD.fna -o final_predicted_viruses+GPD.genes -a final_pre
dicted_viruses+GPD.faa -p meta

##Installing vCONTACT-2
conda create --prefix=vcontact2-0.9.1 -c bioconda vcontact2 mcl blast diamond
cd vcontact2-0.9.19
wget https://paccanarolab.org/static_content/clusterone/cluster_one-1.0.jar

##Clustering the proteins into viral clusters

source activate /scratch/114-canis.familiaris-MINION/condatools/vcontact2-0.9.19
vcontact2_gene2genome -p final_predicted_viruses+GPD.faa -o final_predicted_viruses+GPD_g2g.c
sv -s 'Prodigal-FAA'

vcontact2 --raw-proteins final_predicted_viruses+GPD.faa --rel-mode 'Diamond' --proteins-fp f
inal_predicted_viruses+GPD_g2g.csv --db 'ProkaryoticViralRefSeq201-Merged' --pcs-mode MCL --v
cs-mode ClusterONE --c1-bin /scratch/114-canis.familiaris-MINION/condatools/vcontact2-0.9.19/
cluster_one-1.0.jar -t 40 --output-dir out_viruses_vCONTACT-2_FINAL+GPD


##If it runs out of memory, you can continue/finish the previous command by using this one:

vcontact2 --contigs out_viruses_vCONTACT-2_FINAL+GPD/vConTACT_contigs.csv --pcs out_viruses_v
CONTACT-2_FINAL+GPD/vConTACT_pcs.csv --pc-profiles out_viruses_vCONTACT-2_FINAL+GPD/vConTACT_
profiles.csv --proteins-fp HQ+MQ_final+GPD_g2g.csv --db 'ProkaryoticViralRefSeq201-Merged' --
pcs-mode MCL --vcs-mode ClusterONE --c1-bin /scratch/114-canis.familiaris-MINION/condatools/v
contact2-0.9.19/cluster_one-1.0.jar
```
